# Assessment of different techniques and markers to distinguish recrudescence from new infection in an antimalarial therapeutic efficacy study conducted in Rwanda

**DOI:** 10.1101/2025.08.04.668484

**Authors:** Sara L. Cantoreggi, Monica Golumbeanu, Michaela Zwyer, Aline Uwimana, Jean Damascene Niyonzima, Aimable Mbituyumuremyi, Naomi W. Lucchi, Mateusz M. Plucinski, Christian Nsanzabana

## Abstract

**Introduction:** Accurate estimation of antimalarial drug efficacy against *P. falciparum* requires PCR correction to distinguish recrudescence from new infection in recurrent infections. Genotyping length polymorphic or SNP-rich markers, and various decision algorithms based on match counting or probabilistic approaches may be used for this purpose. In this study, we compared several markers and decision algorithms using samples collected in a therapeutic efficacy study conducted in Rwanda to identify the most suitable and robust approaches for PCR correction.

**Methods:** We optimized nested PCR assays to genotype four microsatellites and assessed their sensitivity in detecting minority clones with laboratory parasite strain mixtures. We analyzed patient samples by capillary electrophoresis and amplicon deep sequencing and assessed the diversity and allelic frequency of *msp1*, *msp2*, *glurp*, microsatellites (*Poly-α, PfPK2, TA40, TA81*), and SNP-rich markers (*ama1-D2, ama1-D3, cpmp, cpp, csp*). We then classified the samples into recrudescence or new infection based on different marker combinations (*msp1/msp2/glurp, msp1/msp2/*microsatellite, and SNP-rich markers) and three decision algorithms (WHO algorithm, 2/3 algorithm, and a Bayesian approach), and compared our results with previously published findings.

**Results:** Among microsatellites, *TA40* and *PfPK2* had the highest sensitivity in detecting minority clones; however, their intra-assay reproducibility was found to be limited. The 3D7 allelic family of the *msp2* gene, *glurp* and SNP-rich markers had the highest genetic diversity. In distinguishing recrudescence from new infection, the WHO algorithm identified the fewest recrudescences across all marker combinations. Conversely, the 2/3 algorithm identified the highest number of recrudescences, and the algorithm based on Bayesian statistics yielded intermediate results. We observed the least variability in classification results among the different algorithms for the SNP-rich markers *ama1-D2/ama1-D3/cpmp* combination. In our study, replacing *glurp* with one of the four analyzed microsatellites or using SNP-rich markers did not significantly alter the final number of recrudescent infections.

**Interpretation:** Our findings emphasize the importance of using assays with high sensitivity in detecting minority clones, and markers with high diversity and low allelic frequency for accurate PCR correction. The SNP-rich markers method yielded the most consistent results in detecting recrudescences, regardless of the decision algorithm used. Therefore, it holds great potential for performing reliable PCR correction. Further evaluation of decision algorithms based on probabilistic approaches, compared to match-counting methods, is essential to ensure the accuracy and consistency of PCR-corrected drug efficacy estimations.

## INTRODUCTION

Gains in malaria control and elimination are threatened by the development of resistance to antimalarial drugs against *Plasmodium falciparum*. Indeed, artemisinin (ART) partial resistance and subsequent selection of resistance to the partner drug in Artemisinin-based Combination Therapies (ACT) is a cause of concern (1, 2). After emerging in Southeast Asia in the late 2000s (3), partial ART resistance quickly developed into full ACT resistance, leading to treatment failures in multiple countries (2, 4). In recent years, ART resistance and mutations associated with partner drug resistance have been increasingly reported in East Africa and the Horn of Africa (5–8), although full ACT resistance has not been reported yet. It is therefore crucial to strengthen drug efficacy monitoring through therapeutic efficacy studies (TESs).

The assessment of antimalarial drug efficacy requires PCR correction to distinguish recrudescence from new infection in recurrent infections (9, 10). Highly polymorphic markers, such as merozoite surface protein 1 (*msp1*), merozoite surface protein 2 (*msp2*), glutamate-rich protein (*glurp*), and multiple microsatellites (11–14) have been used for this purpose. To overcome the limited sensitivity of *glurp* in detecting minority clones in polyclonal infections (15), the World Health Organization (WHO) now recommends to replace this marker with one microsatellite (16). For the transition period, the WHO recommends using both the *msp1*/*msp2*/*glurp* and *msp1*/*msp2*/microsatellite methods for PCR correction to allow for comparison and evaluation (16).

Microsatellites are highly polymorphic (17, 18), but their sensitivity in detecting minority clones has been only partially assessed (19). On the other hand, amplicon deep sequencing (AmpSeq) offers a promising alternative, which has been shown to be highly sensitive in detecting minority clones, reproducible and robust in distinguishing recrudescence from new infection (19, 21). Markers proposed for AmpSeq, such as *cpmp*, *ama1-D2*, *ama1-D3*, *cpp* and *csp*, have shown very high diversity in previous studies (19, 21, 22). However, more data is required to establish a standardized protocol to be applied on a larger scale for TESs.

A decision algorithm allows for classification of sample pairs as recrudescence or new infection (11, 13, 16). Three different decision algorithms have been commonly used in this context: (1) the WHO-recommended (3/3) algorithm, which requires all markers to give concordant results to classify a sample pair as a recrudescence; (2) the 2/3 algorithm, where concordance of two markers is sufficient to determine a recrudescence; and (3) an algorithm based on Bayesian statistics, which calculates the posterior probability (range 0.0 to 1.0) of recrudescence while considering the allelic diversity of each marker in the dataset (23). More recently, an additional decision method based on identity-by-descent (IBD) analysis has been proposed (24, 25).

PCR correction outcomes may vary greatly based on markers and decision algorithms used, especially in high transmission settings where high complexity of infection is common. Therefore, it is essential to use PCR correction methods that accurately distinguish recrudescences from new infections to ensure a precise estimation of drug efficacy. Here, we genotyped samples from a TES conducted in Rwanda in 2018 to evaluate the efficacy of Artemether-Lumefantrine (AL). Seven microsatellites had been previously analyzed with the Bayesian algorithm, and PCR-corrected efficacy of AL was found to be >90% (5). We repeated genotyping with *msp1*/*msp2*/*glurp*, *msp1*/*msp2*/microsatellite (*Poly-α, PfPK2, TA40,* and *TA81*) and five SNP-rich markers (*cpmp, ama1-D2, ama1-D3, cpp* and *csp*). We assessed the genetic diversity of each marker and analyzed the different marker combinations with three decision algorithms, with the aim of comparing markers and algorithms to identify the most robust ones for PCR correction.

## METHODS

### Sample collection

Samples were collected in a TES conducted in Rwanda in 2018 by the Rwanda Biomedical Center (RBC) supported by the United States President’s Malaria Initiative (PMI) and Centers for Disease Control and Prevention (US-CDC). Dried blood spots (DBS) on Whatman 903 filter paper were collected in three sentinel sites: Masaka (Kicukiro district, City of Kigali), Rukara (Kayonza district, Eastern Province) and Bugarama (Rusizi district, Western Province). Detailed information about site characteristics and sample collection has been published elsewhere (5).

### DNA extraction

Three punches of 3 mm diameter were obtained from each DBS. DNA extraction was performed with the QIAamp 96 DNA Blood Kit (Qiagen) according to the manufacturer’s instructions (26).

### Genotyping of length polymorphic markers: *msp1*, *msp2* and *glurp*

*Msp1*, *msp2,* and *glurp*, as well as their respective allelic families (MAD20, K1 and Ro33 for *msp1*, 3D7 and FC27 for *msp2*), were amplified by primary and nested PCR as previously described (15, 19). Amplicon sizes were assessed by high-resolution capillary electrophoresis (H-CE) on ABI 3730xl^®^ (Thermo Fisher Scientific). For Ro33, considered to be monomorphic (27, 28), sample positivity was assessed with the DNA Fast Analysis Kit on QIAxcel (Qiagen). H-CE data was analyzed using GeneMapper v. 6 (Thermo Fisher Scientific). Any peak with fluorescence intensity <500 RFU was considered background noise. Additionally, a cut-off of 10% and 20% on the highest peak’s fluorescence intensity was applied for *msp1*/*msp2* allelic families and *glurp*, respectively, to remove stutter peaks, as indicated in (15).

### Genotyping of length polymorphic markers: microsatellites

A nested PCR assay for *Poly-α*, *PfPK2*, *TA40* and *TA81* was optimized based on previously published protocols (14, 29, 30). Details are available in Supplementary Tables 1-4. Amplicon sizes were assessed by H-CE on ABI 3730xl® (Thermo Fisher Scientific). H-CE data was analyzed using GeneMapper v. 6 (Thermo Fisher Scientific). Any peak with fluorescence intensity <500 RFU was considered background noise. Additionally, a peak-calling algorithm adapted from (22) was used to remove stutter peaks.

### Limit of detection (LOD) for microsatellites

The limit of detection (LOD) of the assay (i.e., its sensitivity in detecting minority clones) was assessed with mixtures of four different laboratory strains of *P. falciparum* (3D7, K1, HB3, and FCB1) in different ratios, as described previously (19). The assay was run in triplicate. We only considered strain mixture samples in which the strain with the longest amplicon was in minority (Supplementary Table 5). A strain mixture sample was classified as positive if the minority clones were detected (i.e., if all strains present in the sample were detected) in two out of three replicates. The LOD was defined as the lowest ratio of laboratory strains at which a strain mixture sample was classified as positive. Intra-assay reproducibility was assessed by calculating the percentage of samples returning the same result for all three replicates.

### Amplicon deep sequencing of SNP-rich markers: *cpmp*, *ama1-D2*, *ama1-D3*, *cpp* and *csp*

*Cpmp* (430 bp), *ama1-D2* (479 bp), *ama1-D3* (516 bp), *cpp* (395 bp), and *csp* (329 bp) were sequenced as described previously (21, 22). Library preparation consisted of primary, nested and adapter PCR, followed by magnetic purification, normalization and sample pooling. Amplicons were then deep sequenced on MiSeq (Illumina) in paired-end mode with the MiSeq reagent kit v3 (600 cycles).

### Bioinformatics analysis of amplicon sequencing data

The R package *HaplotypR* (31) was employed to analyze the amplicon sequencing data and identify haplotypes. Sequencing reads were first demultiplexed by sample and marker. Primer sequences were then removed, and the forward and reverse reads were merged. These merged reads were aligned to the 3D7 *P. falciparum* reference genome, and single nucleotide polymorphisms (SNPs) were identified within each sample. To ensure reliability, only dominant SNPs (occurring at 50% or higher mismatch rate in at least two samples) were considered. Based on these identified SNPs, haplotypes were constructed. Further analysis included only the haplotypes occurring in at least two of the three replicates, having a frequency of 1% or higher, and supported by at least 20 reads.

We further determined the minimum number of reads necessary for a haplotype to be considered true in order to discard spurious haplotypes. The thresholds were determined based on the haplotypes detected in the positive controls. These contained four laboratory strains with known haplotype sequences mixed in different ratios. For most markers, we detected between one and three spurious haplotypes in some controls and hypothesized that these were due to contamination and/or to sequencing errors. To remove them, we applied marker-specific cut-offs. For *ama1-D2,* a haplotype had to be supported by at least 15% of the number of reads of the haplotype with the highest number of reads within a sample. For *ama1-D3*, *cpmp* and *cpp* a cut-off of 25%, 4%, and 8% was defined, respectively. Haplotypes detected in the same sample with number of reads below the threshold were discarded. No threshold was applied for *csp* because no spurious haplotypes were detected in the positive controls. The same thresholds were applied to patient samples.

### Genetic diversity and expected heterozygosity of length polymorphic and SNP-rich markers

Genetic diversity was assessed by counting the number of genotypes (for length polymorphic markers) and haplotypes (for SNP-rich markers), and the frequency at which they were present in the samples from the dataset. Amplicons that were within 3 bp (for *msp1*/*msp2/glurp*) and 2 bp (for microsatellites) of difference were considered identical. Additionally, the expected heterozygosity (H_e_) was calculated for each marker with the following formula: *H*_*e*_ = [*n*/(*n* − 1)][1 − ∑ *p*_*i*_ ^2^], where *n* is the number of genotyped samples and *p_i_* is the frequency of each allele (32).

### Multiplicity of infection analysis

Multiplicity of infection (MOI) for each sample, defined as the number of clones which co-infect a single host, was calculated as the count of all clones detected in the respective sample for each marker. The MOI of the different markers was then stratified by site (Masaka, Rukara, and Bugarama) and used as a proxy for marker diversity, sensitivity and transmission intensity. The Kruskal-Wallis test (33) was used to assess whether the differences between MOI values among all sites or markers were statistically significant. When this test yielded significant results, the post-hoc Dunn’s test with Bonferroni adjustment (34) was used for pairwise comparisons between individual sites or markers. Three markers with the highest MOI values (one from each marker type: *msp1/msp2/glurp*, microsatellites, and SNP-rich markers) were selected for further analysis of MOI in individual day 0 and day X samples. Concordance between two markers was defined as the percentage of samples in which both markers detected the same MOI.

### Classification of infections

Recrudescence vs. new infection analysis was performed by comparing genotypes from the sample collected before treatment (day 0) and from the sample collected in the recurrent infection (day X, where “X” corresponds to the day of recurrence). For length polymorphic markers, the size of the amplicons detected in the day 0 and day X samples was compared. Amplicons that were within 3 bp (for *msp1*/*msp2/glurp*) and 2 bp (for microsatellites) of difference were considered as having the same size. For SNP-rich markers, the haplotypes detected in the day 0 and day X samples were compared. When at least one common haplotype was found on both day 0 and day X, the sample pair was considered a recrudescence. Different marker combinations were used to classify recurrent infections as either recrudescent or new infections, namely *msp1/msp2/glurp*, *msp1/msp2/Poly-α*, *msp1/msp2/TA81, msp1/msp2/TA40*, *msp1/msp2/PfPK2*, *cpmp/ama1-D3/csp*, *cpmp/ama1-D2/csp*, *cpmp/ama1-D3/cpp*, *cpmp/ama1-D2/cpp*, *csp/ama1-D3/cpp*, *csp/ama1-D2/cpp*, *ama1-D2/ama1-D3/csp*, *ama1-D2/ama1-D3/cpmp*, *ama1-D2/ama1-D3/cpp,* and *cpmp/cpp/csp*.

The different decision algorithms, namely the WHO, 2/3 and Bayesian algorithms, were applied to each marker combination. For the Bayesian algorithm we used 100’000 iterations and 2 chains to assess and ensure convergence. The probability of recrudescence for each sample pair was calculated by taking the mean of the probabilities estimated in the 2 chains. Sample pairs with probability of recrudescence >0.5 were considered recrudescent. To calculate the total number of recrudescences within the dataset, all probabilities of recrudescence were summed up. Identity-by-descent analysis was performed for SNP-rich markers with *Dcifer*, a method that allows to infer the degree of shared ancestry between polyclonal infections (25). The corresponding R package *dcifer*, version 1.2.1 (https://cran.r-project.org/web/packages/dcifer/index.html) was used. The default *dcifer* settings were used except for the “lrank” parameter, which was set to 1 given the low number of markers in analysis.

## RESULTS

### Limit of detection of nested PCR assay for microsatellites

The limit of detection (LOD) for minority clones varied among markers and replicates. In general, the LOD was very low for *TA81* and *Poly-α*, as all four laboratory strains were detectable only up to the 5:1:1:1 ratio. For *TA40,* the LOD was three times higher, with all strains detected up to a 1:1:15:1 ratio. *PfPK2* had the highest LOD, detecting all strains at a 1:50:1:50 ratio, about ten times higher than *TA81* and *Poly-α* (Figure 1a). Intra-assay reproducibility was moderately high for *TA40*, *PfPK2* and *Poly-α* (concordance of 63.6%), and lower for *TA81* (concordance of 18.2%) (Figure 1b).

**Figure 1.**
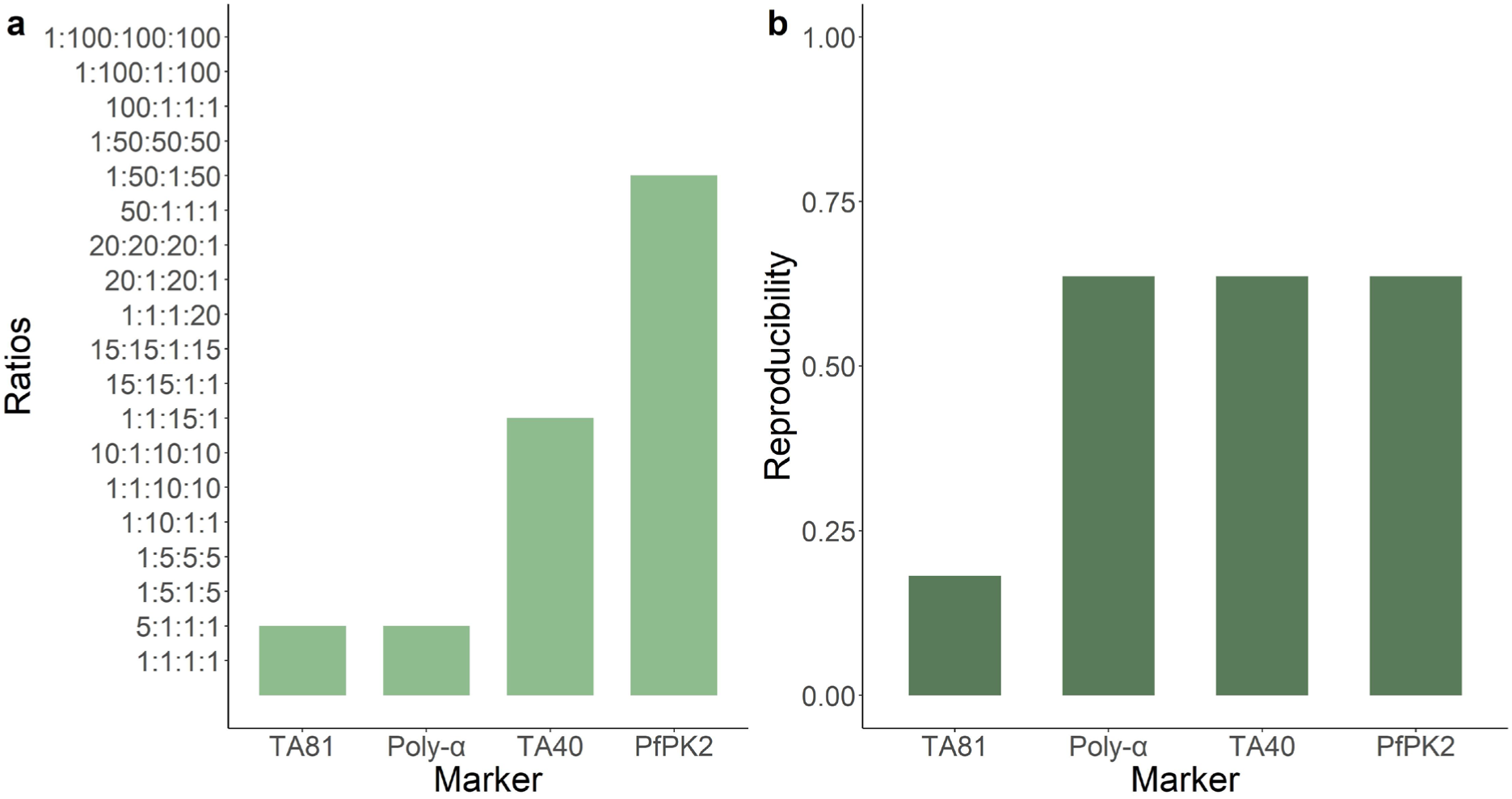
Limit of detection and intra-assay variability of microsatellites *TA81, Poly-α*, *TA40* and *PfPK2*. The assay was tested with four laboratory strains (3D7, K1, HB3, FCB1) mixed in different ratios. (**a**) Limit of detection (LOD) of the assay for each microsatellite (x-axis). Y-axis: ratios of the strain mixture samples. The height of the bars corresponds to the LOD value. (**b**) Intra-assay reproducibility for each microsatellite (x-axis). The height of the bars corresponds to the reproducibility value (0.0 to 1.0, y-axis).

### Diversity and allelic frequency of markers used

A total of 256 patient samples, including 188 unpaired and 34 paired samples, were successfully analyzed for *msp1, msp2, glurp, TA40, TA81, Poly-α,* and *PfPK2*. Additionally, the 34 paired samples were successfully sequenced for SNP-rich markers. The full sample list and a flow chart illustrating the number and origin of samples analyzed are available in Supplementary Table 6 and Supplementary Figure 1.

Genetic diversity and expected heterozygosity (H_e_) varied across the markers (Figure 2, Table 1). When considering *msp1*, *msp2* and *glurp*, *msp2*/3D7 and *glurp* had the highest diversity, with 65 (H_e_ = 0.976) and 39 (H_e_ = 0.954) genotypes detected, respectively. *Msp1*/MAD20, *msp1*/K1 and *msp2*/FC27 were less diverse, with 25 (H_e_ = 0.927), 24 (H_e_ = 0.908), and 33 (H_e_ = 0.883) genotypes, respectively. For *msp2*/FC27, 3 genotypes had high allelic frequencies of 23%, 18% and 15%, and were present in more than half of the isolates. Microsatellites generally exhibited lower diversity than the other length polymorphic markers. *Poly-α* was the most diverse, with 23 distinct genotypes (H_e_ = 0.923), followed closely by *TA40* (23 genotypes, H_e_ = 0.881), and *TA81* (12 genotypes, H_e_ = 0.853). *PfPK2* was the least diverse microsatellite, with one dominant genotype exhibiting an allelic frequency of 34% (16 genotypes, H_e_ = 0.834). All SNP-rich markers had high diversity. *Cpmp* had the highest diversity, with 52 haplotypes (H_e_ = 0.981), followed by *cpp* with 39 haplotypes (H_e_ = 0.961). *Ama1-D2* and *ama1-D3*, two fragments of the *ama1* gene, yielded very similar diversity values, as expected, with each 38 haplotypes and expected heterozygosity, H_e_, of 0.955 and 0.959, respectively. *Csp* also showed similar diversity with 35 haplotypes (H_e_ = 0.958). The H_e_ for SNP-rich markers was higher compared to length polymorphic markers, except *msp2*/3D7.

**Figure 2.**
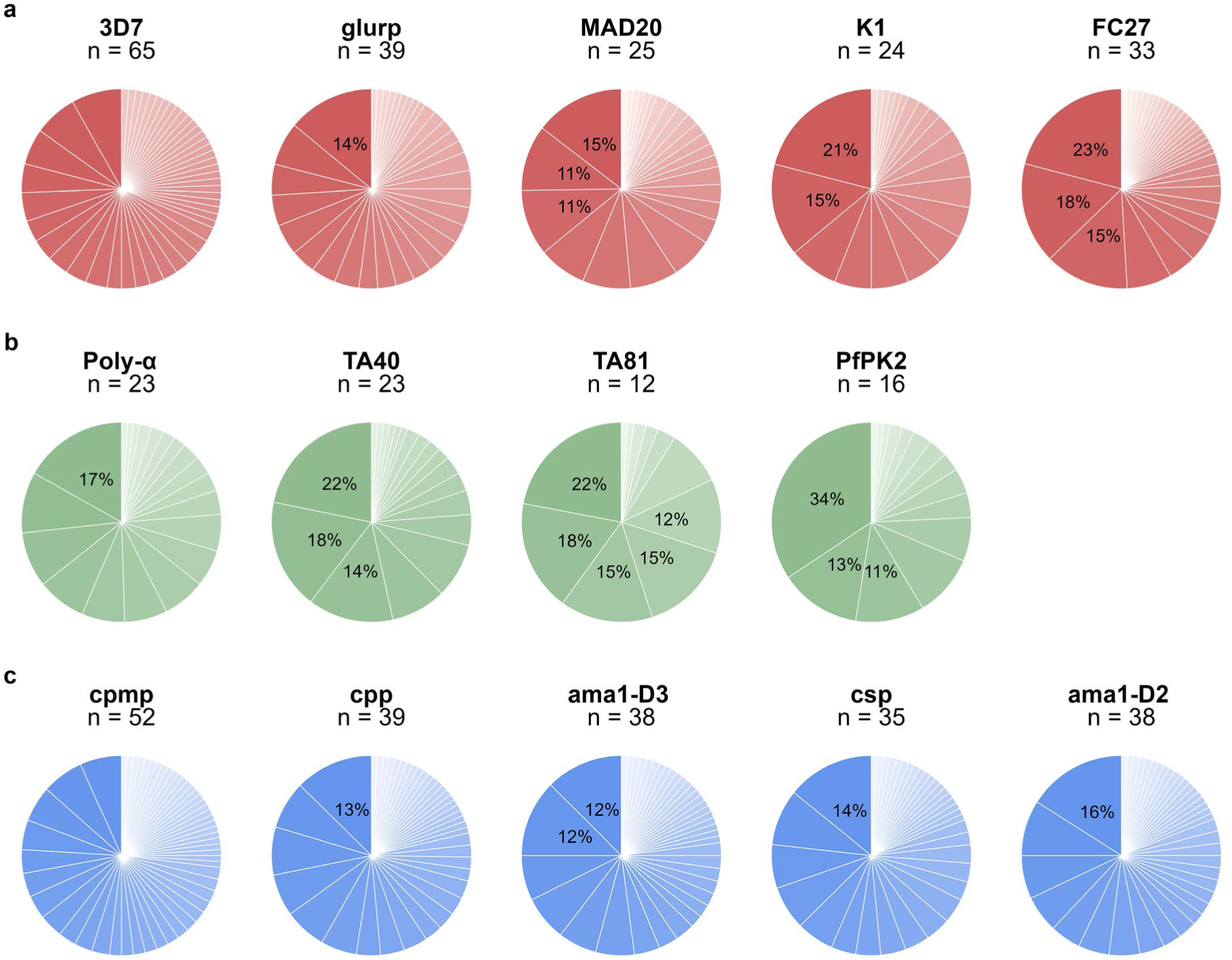
Diversity and allelic frequency of different markers. Each pie chart represents the allelic diversity of one marker. The detected alleles are indicated with different shades of each color, while their frequency is represented by the size of the slice and indicated in percentage (%). Only frequencies > 10% are indicated. (**a**) *msp1* (MAD20, K1), *msp2* (3D7, FC27), and *glurp*. (**b**) Microsatellites (*Poly-α, TA40, TA81*, and *PfPK2*). (**c**) SNP-rich markers (*cpmp, ama1-D2, ama1-D3, cpp*, and *csp*). Each row is ordered by decreasing expected heterozygosity (i.e., decreasing diversity). n = number of genotypes/haplotypes.

**Table 1.**
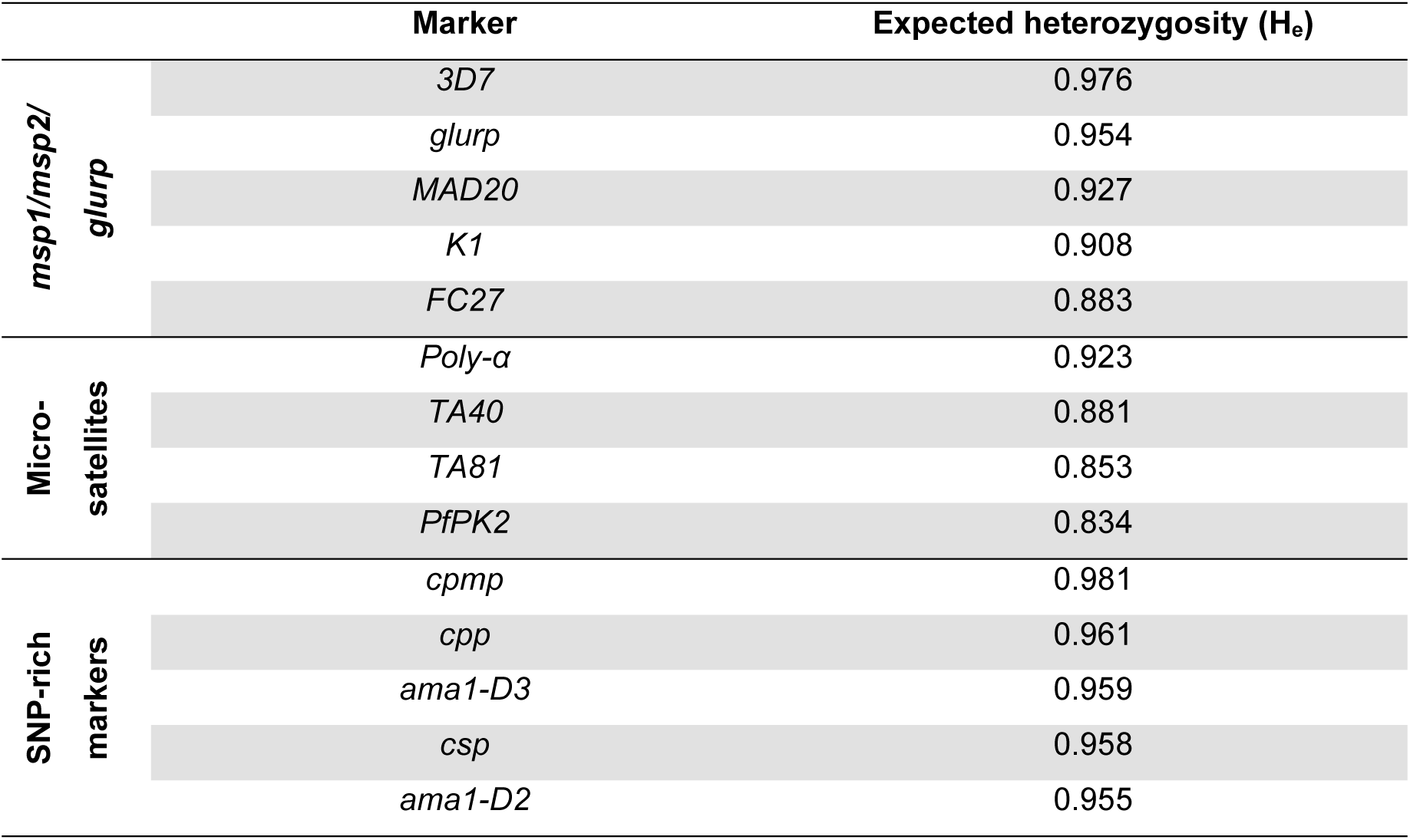
Expected heterozygosity of markers. Values vary between 0 (no diversity) and 1 (high diversity). Markers are listed by decreasing expected heterozygosity.

### Multiplicity of infection

MOI varied among sites and markers (Figure 3). In general, the MOI was slightly lower in Masaka (mean across all markers: 1.47, range 1-4), followed by Bugarama (mean: 1.64, range 1-6), and Rukara (mean: 1.77, range 1-7). Differences in MOI were found to be statistically significant between Bugarama and Masaka (adj. *p-value* = 5.97·10^−3^) and between Rukara and Masaka (adj. *p-value* = 8.18·10^−6^). Among the markers, *msp1* revealed the highest MOI (mean: 2.41, range 1-7), followed by *msp2* (mean: 2.36, range 1-7), *csp* (mean: 1.99, range 1-6)*, cpmp* (mean: 1.82, range 1-6), and *cpp* (mean; 1.60, range 1-5). The lowest MOI values were detected by *ama1-D2* and *D3*, the microsatellites and *glurp*. No statistically significant pairwise differences between markers were found (adj. *p-values* > 0.05). *Msp1*, *Poly-α,* and *csp* had the highest MOI values among their respective marker type and were thus selected for analysis of MOI in the 34 day 0/day X sample pairs (Supplementary Figure 2). For day 0 samples, all three markers detected the same number of clones (i.e., same MOI) in 14.7% (5/34) of the samples. *Poly-α* and *csp* detected the same number of clones in 50% (17/34) of the samples, while *msp1* and *Poly-α* and *msp1* and *csp* had lower concordance rates (23.5% (8/34) and 29.4% (10/34), respectively). For day X samples, the concordance between the three markers was higher. Indeed, all three found the same MOI in 35.3% (12/34) of the samples, and all other marker combinations had similar concordance rates (*msp1* and *Poly-α*: 52.9% (18/34), *msp1* and *csp*: 52.9% (18/34), *Poly-α* and *csp*: 50% (17/34)).

**Figure 3.**
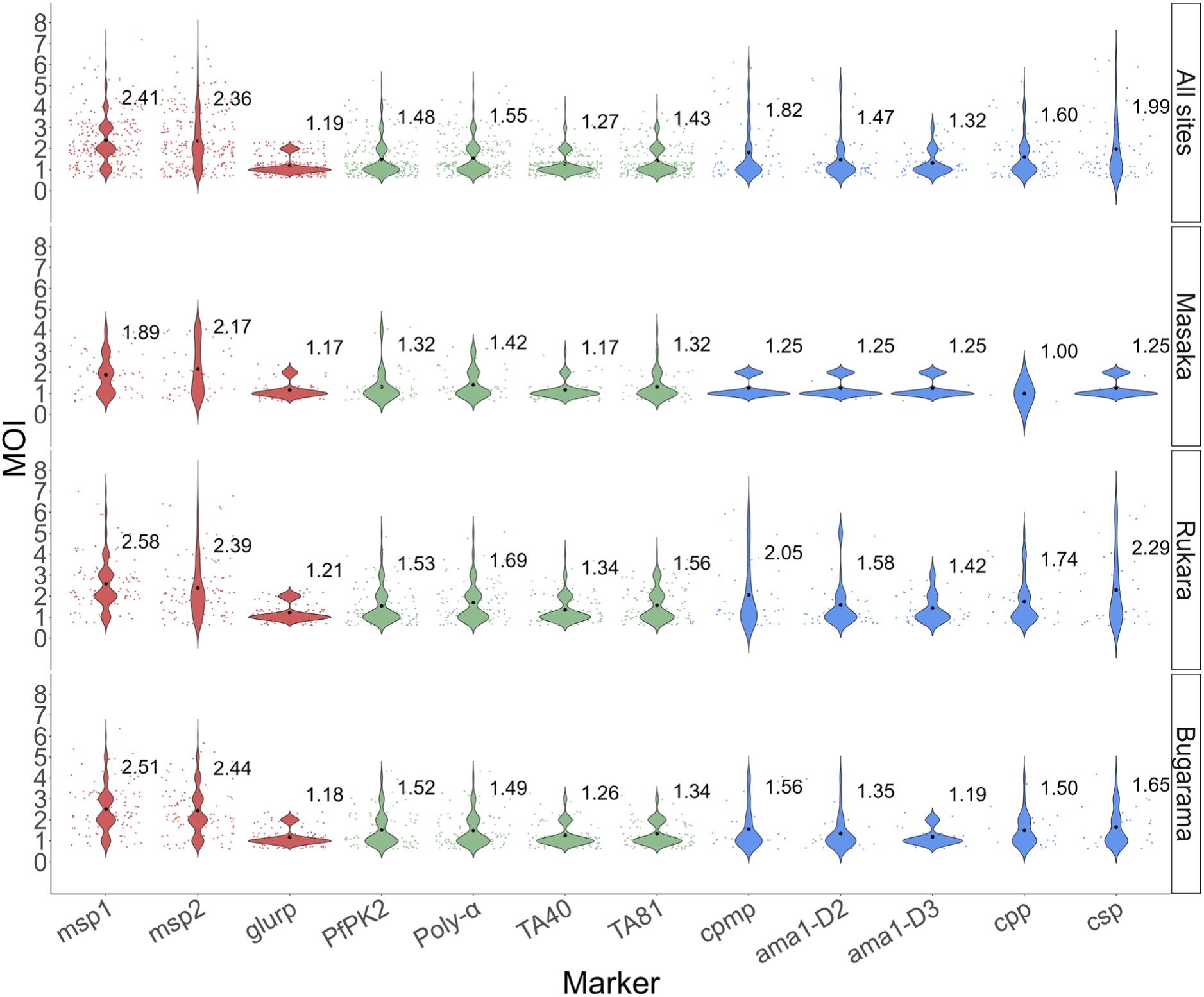
MOI detected by each marker in all sites and stratified by site (Masaka, Rukara, and Bugarama). The violin plots represent the distribution of MOI values. The black dots represent the mean MOI (value indicated next to each violin), while the colored dots represent the MOI of each sample. Markers: *msp1*, *msp2, glurp*, *PfPK2, Poly-α*, *TA40*, *TA81*, *cpmp, ama1-D2, ama1-D3, cpp* and *csp*. For length polymorphic markers, 256 samples were analyzed. For AmpSeq markers, 68 samples were analyzed, and only 4 of them were from Masaka.

### Classification of infections

The total number of recrudescences varied among the different marker combinations and decision algorithms, as shown in Figure 4. One sample pair (01/03) was excluded from the analysis because it was negative for *cpp.* The WHO algorithm classified the least amount of recrudescences compared to the other algorithms. Indeed, it classified on average 6.4 recrudescences (range: 5-9). The 2/3 algorithm classified the highest (mean: 10.5) and least consistent amount of recrudescences, varying from 7 with *ama1-D3/cpmp/cpp* to 19 with *msp1/msp2/TA81* (range: 7-19). The Bayesian algorithm found on average 8.8 recrudescences (range: 6.2-10.7). The three algorithms showed the most consistent results with the SNP-rich markers combinations, in particular with the *ama1-D3/cpmp/cpp*, *ama1-D2/cpmp/cpp, ama1-D2/ama1-D3/csp* and *ama1-D2/ama1-D3/cpmp* combinations (Table 2). The *ama1-D2/ama1-D3/cpmp* combination provided the most consistent results independent of the algorithm used; therefore, we assumed that it would provide the most accurate classification results and used it as the “reference method” for all further analyses.

**Figure 4.**
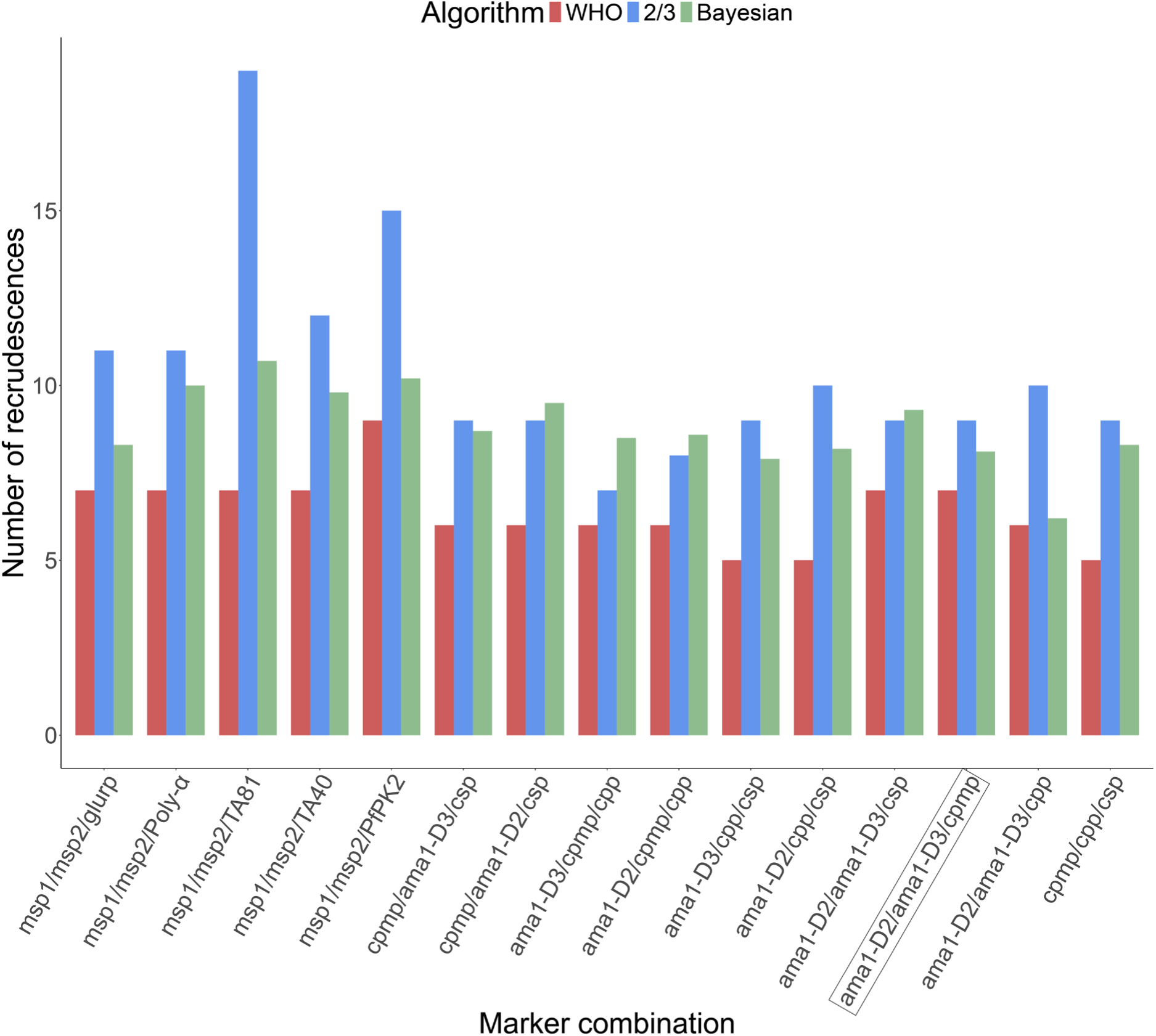
Number of recrudescences found with each marker combination and algorithm. Number of recrudescences is shown on the y-axis. Marker combinations (x-axis): *msp1/msp2/glurp*, *msp1/msp2/Poly-α*, *msp1/msp2/TA81*, *msp1/msp2/TA40*, *msp1/msp2/PfPK2*, *cpmp/ama1-D3/csp, cpmp/ama1-D2/csp, cpmp/ama1-D3/cpp, cpmp/ama1-D2/cpp, csp/ama1-D3/cpp, csp/ama1-D2/cpp, ama1-D2/ama1-D3/csp, ama1-D2/ama1-D3/cpmp* (highlighted, chosen as “reference method”), *ama1-D2/ama1-D3/cpp, and cpmp/cpp/csp*. Classification algorithms: WHO, 2/3, Bayesian. For the Bayesian algorithm, the number of recrudescences corresponds to the sum of all posterior probabilities of recrudescence.

**Table 2.** Number of recrudescences with each algorithm, mean, standard deviation. The *ama1-D2/ama1-D3/cpmp* combination (**bold**) was chosen as “reference method”.

|  | WHO<br>algorithm | 2/3<br>algorithm | Bayesian<br>algorithm | Mean | Standard<br>deviation |
| --- | --- | --- | --- | --- | --- |
| *msp1/msp2/glurp* | 7 | 11 | 8.3 | 8.77 | 1.66 |
| *msp1/msp2/Poly-α* | 7 | 11 | 10 | 9.33 | 1.70 |
| *msp1/msp2/TA81* | 7 | 19 | 10.7 | 12.23 | 5.02 |
| *msp1/msp2/TA40* | 7 | 12 | 9.8 | 9.6 | 2.05 |
| *msp1/msp2/PfPK2* | 9 | 15 | 10.2 | 11.4 | 2.59 |
| *cpmp/ama1-D3/csp* | 6 | 9 | 8.7 | 7.9 | 1.35 |
| *cpmp/ama1-D2/csp* | 6 | 9 | 9.5 | 8.17 | 1.55 |
| *ama1-D2/ama1-D3/csp* | 7 | 9 | 9.3 | 8.43 | 1.02 |
| ***ama1-D2/ama1-D3/cpmp*** | 7 | 9 | 8.1 | 8.04 | 0.82 |
| *cpmp/cpp/csp* | 5 | 9 | 8.3 | 7.43 | 1.75 |
| *ama1-D2/ama1-D3/cpp* | 6 | 10 | 6.2 | 7.4 | 1.84 |
| *ama1-D2/cpmp/cpp* | 6 | 8 | 8.6 | 7.53 | 1.11 |
| *ama1-D2/cpp/csp* | 5 | 10 | 8.2 | 7.73 | 2.07 |
| *ama1-D3/cpmp/cpp* | 6 | 7 | 8.5 | 7.16 | 1.03 |
| *ama1-D3/cpp/csp* | 5 | 9 | 7.9 | 7.3 | 1.69 |

Variations between the marker combinations were largely driven by a subset of samples, as shown in Figure 5. Indeed, 81.8% (27/33), 54.5% (18/33) and 72.7% (24/33) of the samples were assigned the same classification (recrudescence (R), in red, or new infection (NI), in blue) with all marker combinations by the WHO, 2/3, and Bayesian algorithm, respectively. The remaining samples were classified differently depending on the marker combination. Concordance between the different algorithms was observed in 39.4% (13/33) of the samples independently of the marker combination used.

**Figure 5.**
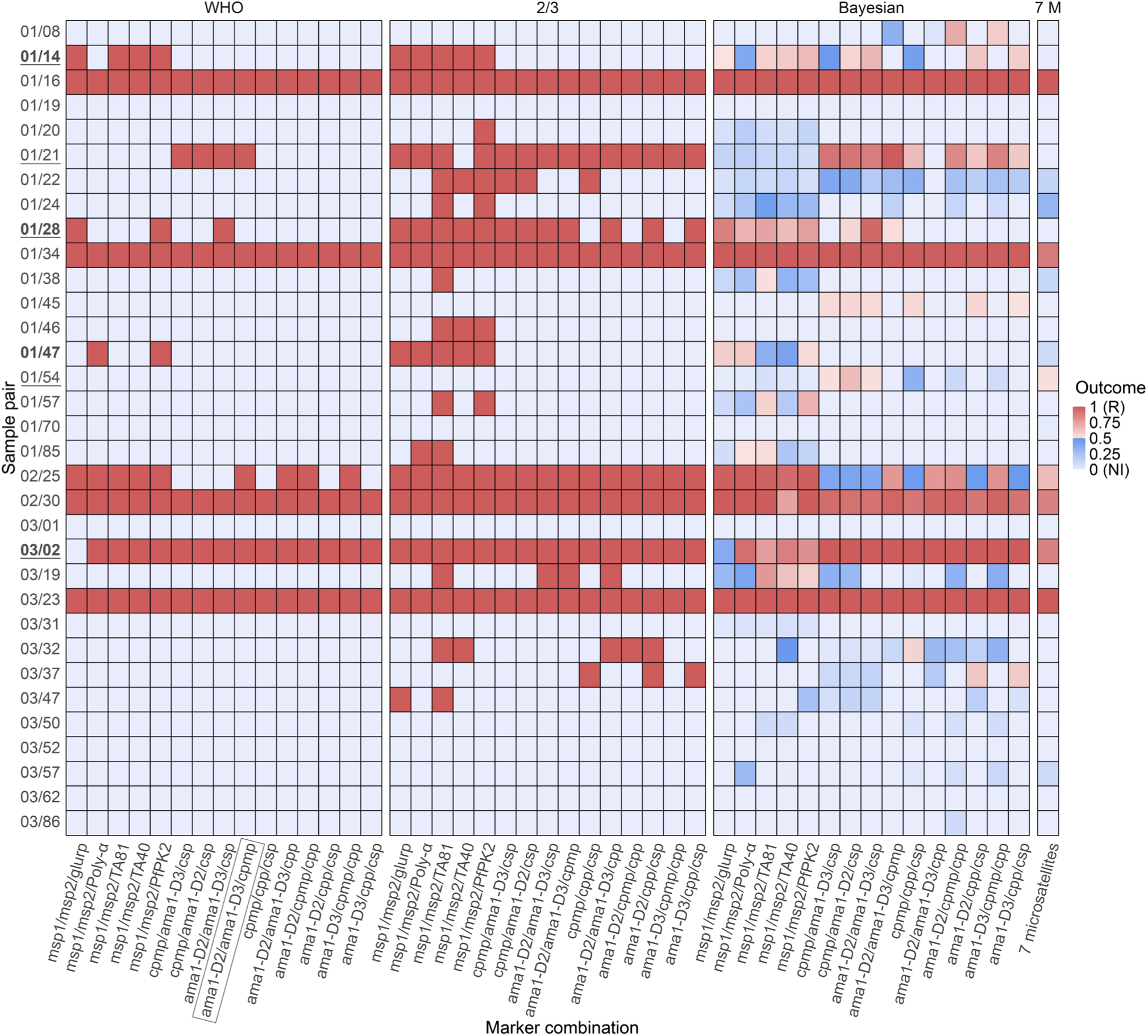
Recrudescence vs. new infection classification of all sample pairs with the different algorithms. Sample IDs are indicated on the y-axis. Marker combinations (x-axis): *msp1/msp2/glurp*, *msp1/msp2/Poly-α*, *msp1/msp2/TA81*, *msp1/msp2/TA40*, *msp1/msp2/PfPK2*, *cpmp/ama1-D3/csp, cpmp/ama1-D2/csp, cpmp/ama1-D3/cpp, cpmp/ama1-D2/cpp, csp/ama1-D3/cpp, csp/ama1-D2/cpp, ama1-D2/ama1-D3/csp, ama1-D2/ama1-D3/cpmp* (highlighted, chosen as “reference method”), *ama1-D2/ama1-D3/cpp, and cpmp/cpp/csp*. Classification algorithms: WHO, 2/3, Bayesian. Previously published data from (5) was added as “7 M” (for 7 microsatellites). Outcomes are colored in a gradient from light blue (prob. of R: 0) to dark blue (prob. of R: 0.49) for new infections, and from light pink (prob. of R: 0.5) to dark red (prob. of R: 1) for recrudescences. **Bold**: samples which showed discordant R vs. NI results when replacing *glurp* with a microsatellite. <u>Underlined</u>: samples which showed discordant R vs. NI results depending on the method used (*msp1/msp2/glurp* with WHO algorithm, *ama1-D2/ama1-D3/cpmp* with WHO algorithm, 7 microsatellites with Bayesian algorithm (5)).

When replacing *glurp* with each of the four microsatellites and applying the WHO algorithm, the same number of recrudescences (7) was found, except when using the microsatellite *PfPK2* which detected 9 recrudescences. Interestingly, the recrudescences were not found in the same sample pairs for all marker combinations. For 87.9% (29/33) of the sample pairs the results were concordant, but 4 sample pairs gave discordant results depending on the marker combination. The discordant cases are indicated in **bold** in Figure 5 (results by marker combination) and in Supplementary Table 7 (results by single marker).

### Recrudescence vs. new infection: comparison with previous results

The results obtained during this study were compared to those of a study published previously (5), which employed the 7 microsatellites assay analyzed with the Bayesian algorithm approach. With the “reference method” *ama1-D2/ama1-D3/cpmp* analyzed with the WHO algorithm, 7 recrudescences were detected. The *msp1/msp2/glurp* combination with the WHO algorithm and the 7 microsatellites analyzed with the Bayesian algorithm also detected 7 and 6.54 (7, rounded) recrudescences, respectively. In this case, too, recrudescences were not found in the same sample pairs. For 84.4% (28/33) of the sample pairs, the results were concordant between the different methods used, but 5 pairs gave discordant results, as shown <u>underlined</u> in Figure 5 (results by marker combination) and in Supplementary Table 8 (results by single marker).

### Identity-by-descent analysis

Identity-by-descent analysis of paired samples was conducted on SNP-rich markers data and revealed statistically significant high genetic relatedness (IBD = 1) for 5/33 sample pairs (Table 3). All these pairs were also classified as a recrudescence by the WHO and 2/3 algorithms and had a Bayesian posterior probability of recrudescence >0.962 for the *ama1-D2/ama1-D3/cpmp* combination. Two other sample pairs (01/21 and 02/35) had statistically significant moderate genetic relatedness (IBD >0.5, where a 0.5 value is expected for full siblings (35)). Both were classified as recrudescence by the WHO, 2/3, and the Bayesian algorithms (Table 3).

**Table 3.** Comparison of IBD values with WHO and Bayesian classification algorithms. IBD analysis was performed on *cpmp, ama1-D2, ama1-D3, csp*, and *cpp.* Results are compared to those obtained with WHO, 2/3, and Bayesian algorithms for the *ama1-D2/ama1-D3/cpmp* combination. For the Bayesian algorithm, a sample pair is considered a recrudescence when the posterior probability of recrudescence is >0.5. R: recrudescence, NI: new infection.

| Sample pair | IBD | *p*-value (IBD) | WHO and 2/3 algorithm | Bayesian algorithm |
| --- | --- | --- | --- | --- |
| 01/16D0<br>01/16D21 | 1 | $2.67 \cdot 10^{-8}$ | R | 1.000 |
| 01/21D0<br>01/21D21 | 0.665 | $2.58 \cdot 10^{-3}$ | R | 0.996 |
| 01/34D0<br>01/34D21 | 1 | $6.74 \cdot 10^{-6}$ | R | 0.989 |
| 02/25D0<br>02/25D28 | 0.778 | $5.82 \cdot 10^{-5}$ | R | 0.775 |
| 02/30D0<br>02/30D21 | 1 | $2.42 \cdot 10^{-8}$ | R | 0.962 |
| 03/02D0<br>03/02D21 | 1 | $1.38 \cdot 10^{-7}$ | R | 0.999 |
| 03/23D0<br>03/23D21 | 1 | $1.17 \cdot 10^{-7}$ | R | 0.999 |

## DISCUSSION

Accurately estimating antimalarial drug efficacy through therapeutic efficacy studies (TESs) is crucial to inform policy makers, especially in regions where artemisinin partial resistance has emerged. An overestimation of drug efficacy could lead to delays in recognizing the spread and magnitude of drug resistance, and prevent necessary action to mitigate the impact of reduced drug efficacy, while an underestimation of efficacy could lead to premature changes in treatment policies, possibly leading to lower access to treatment for the transition period and additional costs for malaria programs (16). Accurate estimation of drug efficacy requires robust and reproducible PCR correction methods which utilize highly sensitive assays and markers with high genetic diversity and low allelic frequencies. Additionally, it requires a decision algorithm which allows for correct classification of a recurrent infection as a recrudescence or a new infection based on several markers.

Previous studies have shown that SNP-rich markers analyzed with AmpSeq and *msp1*/*msp2* analyzed with high-resolution capillary electrophoresis (H-CE) have very high sensitivity in detecting minority clones, while *glurp* has much lower sensitivity (19, 21). We had initially assessed the sensitivity of microsatellites in detecting minority clones using single-round PCR and found it to be moderate (19). Here, we aimed to assess the sensitivity of a nested PCR assay for four microsatellites, expecting higher sensitivity compared to the single-round PCR. Out of the four microsatellites tested, high sensitivity in detecting minority clones was observed for *TA40* and *PfPK2*. For the latter, the sensitivity was comparable to that of the AmpSeq assay, detecting minority clones in a 1:50:1:50 ratio. However, high variability between replicates was observed, in contrast to the one-round PCR assay, which was previously shown to be highly reproducible with low intra- and inter-assay variability (19, 36).

Low marker diversity and high allelic frequency can lead to an overestimation of recrudescences and thus an underestimation of drug efficacy. Therefore, it is critical to use highly diverse markers with low allelic frequencies. Here, we report high diversity (H_e_ > 0.95) and low allelic frequency for *msp2/3D7*, *glurp* and all SNP-rich markers, and lower diversity (H_e_ < 0.90) for *msp2/FC27* and most of the microsatellites, with genotypes harboring higher allelic frequencies, thus possibly introducing a bias towards a recrudescence classification for non-statistical methods like match counting approaches that do not take allele frequency into account. This is in line with previous studies, where *msp2/3D7* (19, 28, 37–39), *Poly-α* (40–43), and *cpmp* (19, 21, 22) have been consistently reported to be the most diverse length polymorphic, microsatellite and SNP-rich marker, respectively.

MOI data can be used as a proxy for marker diversity and sensitivity, also reflecting transmission intensity in the study areas. Indeed, we found significant differences in MOI between sites, likely reflecting differences in transmission intensity. As expected, MOI was lower in Masaka, where in 2018 malaria incidence was reported to be 100-250 cases per 1000 population, compared to Bugarama and Rukara, where incidence was > 450 cases per 1000 population (44, 45). Higher MOI values correspond to higher numbers of clones detected in a sample and are therefore associated with higher diversity (different clones have different genotypes), higher sensitivity (minority clones are detected) and higher transmission intensity (multiple clones infect the same individual). Within this dataset, *msp1*, *msp2* and most SNP-rich markers detected high MOI values, confirming their high discriminatory power. The multi-allelic family nature of *msp1* and *msp2* also contributed to higher MOI values for these markers. In contrast, *glurp* and microsatellites generally had lower MOI values, reflecting the low sensitivity and/or low diversity of these markers. *Ama1-D2* and *ama1-D3* also had lower MOI values, but these were likely attributable to the higher thresholds applied to the reads to discard spurious haplotypes, rather than to low sensitivity and diversity.

The WHO recommends to replace *glurp* with the microsatellite with greatest diversity in the study region (16). Our diversity and MOI data suggest that *Poly-α, PfPK2 and TA40* may be adequate candidates for this purpose. To evaluate this hypothesis, we replaced *glurp* with each microsatellite and applied the different decision algorithms to all marker combinations. With the WHO algorithm, we found a consistent and lowest number of recrudescences across marker combinations. This approach requires concordant results from all markers to confirm a recrudescence, and it is therefore expected that a low number of them is detected. The 2/3 algorithm classified the highest number of recrudescences, up to 19 in the case of the *msp1/msp2/TA81* combination. This algorithm was initially introduced to overcome the limitation of *glurp* in detecting minority clones and thus in detecting recrudescent infections (16), however, it likely overestimated recrudescences in this case. The Bayesian algorithm detected a higher number of recrudescences than the WHO algorithm, but a lower number than the 2/3 algorithm, for most marker combinations. By providing a posterior probability of recrudescence based on marker diversity, this algorithm may overcome the limitation of match counting algorithms underestimating or overestimating the number of recrudescences. Based on these results, it is not possible to conclude which algorithm is most accurate; however, probabilistic approaches may have the potential to provide more accurate outcomes compared to match counting approaches (23, 46). Indeed, identity-by-descent (IBD) analysis confirmed significant genetic relatedness of clones in 7 sample pairs and yielded the same results as the Bayesian algorithm.

The number of recrudescences detected was most consistent across algorithms for the *ama1-D2/ama1-D3/cpmp* combination. Due to the minimal variability given by the different algorithms, we argue that AmpSeq with this marker combination may provide the most accurate genotyping results in this context. The comparison of our results with those published previously (5) showed that the *msp1/mp2/glurp* and *ama1-D2/ama1-D3/cpmp* analyzed with the WHO algorithm and the 7 microsatellites analyzed with the Bayesian algorithm gave the same results in terms of number of recrudescences detected, but results were discordant for some sample pairs. Discordant results were due to the low genetic diversity and high allelic frequency of some markers, and/or to the low sensitivity in detecting minority clones of other markers.

Our study has some limitations. First, the nested PCR assay for microsatellites had low reproducibility, which could have resulted in inaccurate results. We also assessed intra-assay variability but did not assess inter-assay variability. Second, erroneously calling stutter peaks (for length polymorphic markers) or haplotypes with low coverage (for SNP-rich markers) as real clones may have led to wrong estimations of diversity, MOI and recrudescences. This is particularly true for microsatellites, for which data analysis is complex due to the high presence of stutter peaks and small size differences between real clones. Third, only 34 sample pairs were used for the analysis of SNP-rich markers, and only 2 of them were collected in Masaka. A bigger sample set would have likely resulted in more generalizable results. Fourth, we found that the *ama1-D2/ama1-D3/cpmp* combination gave the most consistent results and used it as “reference method”. However, *ama1-D2* and *ama1-D3* are two fragments of the same gene (*ama1*), which could make them more likely to be similar in terms of genetic diversity, and thus lead to biased results. Fifth, in order to discard the spurious haplotypes detected in the positive controls, which were likely due to contaminations and/or sequencing errors, we applied quite stringent thresholds, especially for *ama1-D2* and *ama1-D3*. This might have led to the potential loss of true haplotypes. Finally, the generalizability of our results may be limited to settings with similar malaria transmission intensity to Rwanda. Indeed, the relatively low discordance between marker combinations and algorithms reported here may be largely due to low transmission levels and the resulting low complexity of infections in this dataset. In settings with higher transmission, discrepancies between methods may be much more significant, as reported by a recent study conducted in Uganda (42).

The AmpSeq assay has previously proven to be highly sensitive and robust (19) and therefore holds great potential for accurate PCR-corrected drug efficacy estimations. However, the importance of bioinformatics and data analysis steps should not be underestimated. Indeed, applying different thresholds at different analysis steps, such as for example the mismatch rate for SNPs calling or the marker-specific threshold to discard spurious haplotypes, may significantly impact the final genotyping outcome. A careful evaluation and standardization of the pipelines and downstream analysis steps would be necessary to produce robust results and allow for comparison between studies conducted across different locations and time periods.

In conclusion, our results emphasize the importance of using assays with high sensitivity in detecting minority clones, and markers with high diversity and low allelic frequencies, for accurate PCR correction. Both sensitivity and diversity must be considered when choosing a microsatellite to replace *glurp*. In our study, replacing *glurp* with a microsatellite did not significantly change the final number of recrudescent infections, and therefore would not have affected drug efficacy estimations. This was likely due to the low complexity of infections resulting from low transmission intensity in the region. Nonetheless, the SNP-rich markers method yielded the most consistent results in detecting recrudescences, regardless of the decision algorithm used, making it a robust alternative to the currently used length polymorphic markers. Decision algorithms based on probabilistic approaches or genetic relatedness rather than match counting need to be further assessed, as they may provide more accurate efficacy estimations.

## ETHICS

Written informed consent was obtained from all participants as indicated in (5). Samples were anonymous and only used for study purposes.

## CODE AVAILABILITY

All codes used in this analysis will be made available on GitHub.

## ACKNOWLEDGEMENTS

We would like to thank the Rwanda Biomedical Centre (Rwanda) for sharing the patient samples used for this study. We would also like to thank the US President’s Malaria Initiative (Rwanda/USA) and the Centers for Disease Control and Prevention (USA) for sharing their protocol for microsatellites analysis and the PET-PCR results obtained from previous analyses of the samples. H-CE was performed at Microsynth AG (Switzerland), amplicon sequencing was done at the Department of Biosystems Science and Engineering ETH (D-BSSE, Switzerland) and bioinformatics analyses were performed at the Scientific Computing Center (sciCORE) at the University of Basel (Switzerland).

## DISCLAIMER

The findings and conclusions in this report are those of the author(s) and do not necessarily represent the official position of the Centers for Disease Control and Prevention.

## Contributions

S.L.C.: methods development, laboratory analysis, data analysis, visualization, writing (original draft). M.G. and M.Z.: data analysis, writing (review and editing). A.U., J.D.N, A.M., and N.W.L.: sample procurement (2018 TES). M.M.P.: data analysis, writing (review and editing). C.N.: conceptualization, supervision, funding acquisition, resources, writing (review and editing). All authors reviewed and approved the final version of this manuscript.

## REFERENCES

1. Menard D, Dondorp A. Antimalarial Drug Resistance: A Threat to Malaria Elimination. Cold Spring Harb Perspect Med. 2017;7(7).

2. Saunders DL, Vanachayangkul P, Lon C. Dihydroartemisinin-piperaquine failure in Cambodia. N Engl J Med. 2014;371(5):484–5.

3. Dondorp AM, Nosten F, Yi P, Das D, Phyo AP, Tarning J, et al. Artemisinin resistance in Plasmodium falciparum malaria. N Engl J Med. 2009;361(5):455–67.

4. Carrara VI, Lwin KM, Phyo AP, Ashley E, Wiladphaingern J, Sriprawat K, et al. Malaria burden and artemisinin resistance in the mobile and migrant population on the Thai-Myanmar border, 1999-2011: an observational study. PLoS Med. 2013;10(3):e1001398.

5. Uwimana A, Umulisa N, Venkatesan M, Svigel SS, Zhou Z, Munyaneza T, et al. Association of Plasmodium falciparum kelch13 R561H genotypes with delayed parasite clearance in Rwanda: an open-label, single-arm, multicentre, therapeutic efficacy study. Lancet Infect Dis. 2021;21(8):1120–8.

6. Asua V, Conrad MD, Aydemir O, Duvalsaint M, Legac J, Duarte E, et al. Changing Prevalence of Potential Mediators of Aminoquinoline, Antifolate, and Artemisinin Resistance Across Uganda. J Infect Dis. 2021;223(6):985–94.

7. Mihreteab S, Platon L, Berhane A, Stokes BH, Warsame M, Campagne P, et al. Increasing Prevalence of Artemisinin-Resistant HRP2-Negative Malaria in Eritrea. N Engl J Med. 2023;389(13):1191–202.

8. Ishengoma DS, Mandara CI, Bakari C, Fola AA, Madebe RA, Seth MD, et al. Evidence of artemisinin partial resistance in northwestern Tanzania: clinical and molecular markers of resistance. Lancet Infect Dis. 2024;24(11):1225–33.

9. Laufer MK. Monitoring antimalarial drug efficacy: current challenges. Curr Infect Dis Rep. 2009;11(1):59–65.

10. Vestergaard LS, Ringwald P. Responding to the challenge of antimalarial drug resistance by routine monitoring to update national malaria treatment policies. Am J Trop Med Hyg. 2007;77(6 Suppl):153–9.

11. Medicines for Malaria Venture & World Health Organization. Methods and techniques for clinical trials on antimalarial drug efficacy: genotyping to identify parasite populations: informal consultation organized by the Medicines for Malaria Venture and cosponsored by the World Health Organization, 29-31 May 2007, Amsterdam, The Netherlands. https://apps.who.int/iris/handle/10665/43824 2008.

12. World Health Organization. Methods for surveillance of antimalarial drug efficacy. https://www.who.int/docs/default-source/documents/publications/gmp/methods-for-surveillance-of-antimalarial-drug-efficacy.pdf?sfvrsn=29076702_2; 2009.

13. Snounou G, Beck HP. The use of PCR genotyping in the assessment of recrudescence or reinfection after antimalarial drug treatment. Parasitol Today. 1998;14(11):462–7.

14. Centers for Disease Control and Prevention (CDC) - Malaria Branch - Division of Parasitic Diseases and Malaria. Neutral Microsatellite Analysis. 2021.

15. Messerli C, Hofmann NE, Beck HP, Felger I. Critical Evaluation of Molecular Monitoring in Malaria Drug Efficacy Trials and Pitfalls of Length-Polymorphic Markers. Antimicrob Agents Chemother. 2017;61(1).

16. World Health Organization. Informal consultation on methodology to distinguish reinfection from recrudescence in high malaria transmission areas: report of a virtual meeting, 17–18 May 2021. Geneva: World Health Organization; 2021. Licence: CC BY-NC-SA 3.0 IGO. 2021.

17. Zhong D, Koepfli C, Cui L, Yan G. Molecular approaches to determine the multiplicity of Plasmodium infections. Malar J. 2018;17(1):172.

18. Su X, Wellems TE. Toward a high-resolution Plasmodium falciparum linkage map: polymorphic markers from hundreds of simple sequence repeats. Genomics. 1996;33(3):430–44.

19. Schnoz A, Beuret C, Concu M, Hosch S, Rutaihwa LK, Golumbeanu M, et al. Genotyping methods to distinguish Plasmodium falciparum recrudescence from new infection for the assessment of antimalarial drug efficacy: an observational, single-centre, comparison study. Lancet Microbe. 2024:100914.

20. Walsh PS, Fildes NJ, Reynolds R. Sequence analysis and characterization of stutter products at the tetranucleotide repeat locus vWA. Nucleic Acids Res. 1996;24(14):2807–12.

21. Gruenberg M, Lerch A, Beck HP, Felger I. Amplicon deep sequencing improves Plasmodium falciparum genotyping in clinical trials of antimalarial drugs. Sci Rep. 2019;9(1):17790.

22. Lerch A, Koepfli C, Hofmann NE, Kattenberg JH, Rosanas-Urgell A, Betuela I, et al. Longitudinal tracking and quantification of individual Plasmodium falciparum clones in complex infections. Sci Rep. 2019;9(1):3333.

23. Plucinski MM, Morton L, Bushman M, Dimbu PR, Udhayakumar V. Robust Algorithm for Systematic Classification of Malaria Late Treatment Failures as Recrudescence or Reinfection Using Microsatellite Genotyping. Antimicrob Agents Chemother. 2015;59(10):6096–100.

24. Holzschuh A, Lerch A, Nsanzabana C. Multiplexed nanopore amplicon sequencing to distinguish recrudescence from new infection in antimalarial drug trials. bioRxiv. 2024:2024.09.11.612449.

25. Gerlovina I, Gerlovin B, Rodríguez-Barraquer I, Greenhouse B. Dcifer: an IBD-based method to calculate genetic distance between polyclonal infections. Genetics. 2022;222(2).

26. Qiagen. QIAGEN Supplementary Protocol: Isolation of genomic DNA from dried blood spots using the QIAamp 96 DNA Blood Kit 2010 [

27. Noranate N, Prugnolle F, Jouin H, Tall A, Marrama L, Sokhna C, et al. Population diversity and antibody selective pressure to Plasmodium falciparum MSP1 block2 locus in an African malaria-endemic setting. BMC Microbiol. 2009;9:219.

28. Mwingira F, Nkwengulila G, Schoepflin S, Sumari D, Beck HP, Snounou G, et al. Plasmodium falciparum msp1, msp2 and glurp allele frequency and diversity in sub-Saharan Africa. Malar J. 2011;10:79.

29. University of Maryland School of Medicine - Center for Vaccine Development - Malaria Group. Protocol for microsatellite genotyping by unlinked markers. 2012.

30. Anderson TJ, Su XZ, Bockarie M, Lagog M, Day KP. Twelve microsatellite markers for characterization of Plasmodium falciparum from finger-prick blood samples. Parasitology. 1999;119 (Pt 2):113–25.

31. Lerch A, Koepfli C, Hofmann NE, Messerli C, Wilcox S, Kattenberg JH, et al. Development of amplicon deep sequencing markers and data analysis pipeline for genotyping multi-clonal malaria infections. BMC genomics. 2017;18:1–13.

32. Nei M. Estimation of average heterozygosity and genetic distance from a small number of individuals. Genetics. 1978;89(3):583–90.

33. Kruskal-Wallis Test. The Concise Encyclopedia of Statistics. New York, NY: Springer New York; 2008. p. 288–90.

34. Dinno A. Nonparametric Pairwise Multiple Comparisons in Independent Groups using Dunn’s Test. The Stata Journal. 2015;15(1):292–300.

35. Wong W, Wenger EA, Hartl DL, Wirth DF. Modeling the genetic relatedness of Plasmodium falciparum parasites following meiotic recombination and cotransmission. PLoS Comput Biol. 2018;14(1):e1005923.

36. Greenhouse B, Myrick A, Dokomajilar C, Woo JM, Carlson EJ, Rosenthal PJ, et al. Validation of microsatellite markers for use in genotyping polyclonal Plasmodium falciparum infections. Am J Trop Med Hyg. 2006;75(5):836–42.

37. Greenhouse B, Dokomajilar C, Hubbard A, Rosenthal PJ, Dorsey G. Impact of transmission intensity on the accuracy of genotyping to distinguish recrudescence from new infection in antimalarial clinical trials. Antimicrob Agents Chemother. 2007;51(9):3096–103.

38. Somé AF, Bazié T, Zongo I, Yerbanga RS, Nikiéma F, Neya C, et al. Plasmodium falciparum msp1 and msp2 genetic diversity and allele frequencies in parasites isolated from symptomatic malaria patients in Bobo-Dioulasso, Burkina Faso. Parasit Vectors. 2018;11(1):323.

39. Kateera F, Nsobya SL, Tukwasibwe S, Hakizimana E, Mutesa L, Mens PF, et al. Molecular surveillance of Plasmodium falciparum drug resistance markers reveals partial recovery of chloroquine susceptibility but sustained sulfadoxine-pyrimethamine resistance at two sites of different malaria transmission intensities in Rwanda. Acta Trop. 2016;164:329–36.

40. Ishengoma DS, Mandara CI, Madebe RA, Warsame M, Ngasala B, Kabanywanyi AM, et al. Microsatellites reveal high polymorphism and high potential for use in anti-malarial efficacy studies in areas with different transmission intensities in mainland Tanzania. Malar J. 2024;23(1):79.

41. Agaba BB, Anderson K, Gresty K, Prosser C, Smith D, Nankabirwa JI, et al. Genetic diversity and genetic relatedness in Plasmodium falciparum parasite population in individuals with uncomplicated malaria based on microsatellite typing in Eastern and Western regions of Uganda, 2019-2020. Malar J. 2021;20(1):242.

42. Mwesigwa A, Golumbeanu M, Jones S, Cantoreggi SL, Musinguzi B, Nankabirwa JI, et al. Assessment of different genotyping markers and algorithms for distinguishing Plasmodium falciparum recrudescence from reinfection in Uganda. Sci Rep. 2025;15(1):4375.

43. Zhou J, Malaria Branch - Centers for Disease Control and Prevention. Microsatellite Diversity Analysis (personal communication). 2022.

44. U.S. President’s Malaria Initiative (Rwanda). Rwanda Malaria Operational Plan FY 2020. Retrieved from (www.pmi.gov); 2020.

45. Rubuga FK, Moraga P, Ahmed A, Siddig E, Remera E, Moirano G, et al. Spatio-temporal dynamics of malaria in Rwanda between 2012 and 2022: a demography-specific analysis. Infect Dis Poverty. 2024;13(1):67.

46. Plucinski MM, Barratt JLN. Nonparametric Binary Classification to Distinguish Closely Related versus Unrelated Plasmodium falciparum Parasites. Am J Trop Med Hyg. 2021;104(5):1830–5.

