## Supplementary Files for "Assessment of different techniques and markers to distinguish recrudescence from new infection in an antimalarial therapeutic efficacy study conducted in Rwanda"

**Supplementary Table 1. Reaction conditions for *PfPK2* and *Poly-α* (singleplex).**

| | Primary PCR | | Nested PCR | |
|---|---|---|---|---|
| **Reagent** | **Final concentration** | **Volume per reaction** | **Final concentration** | **Volume per reaction** |
| $ddH_2O$ | - | 14.75 µL | - | 16.25 µL |
| Buffer B2 | 1x | 2.5 µL | 1x | 2.5 µL |
| dNTPs | 0.2 mM | 2.5 µL | 0.2 mM | 2.5 µL |
| $MgCl_2$ | 1.5 mM | 1.5 µL | 1.5 mM | 1.5 µL |
| Primers fw+rv | 400 nM | 1 µL | 400 nM | 1 µL |
| HOT FIREPol DNA polymerase ® | 0.05 U/µL | 0.25 µL | 0.05 U/µL | 0.25 µL |
| DNA | - | 2.5 µL | - | 1 µL |
| **Total volume** | - | 25 µL | - | 25 µL |

*Primary PCR products are diluted 1:10 in $ddH_2O$ before being added to nested PCR reaction.

**Supplementary Table 2. Reaction conditions for *TA40* and *TA81* (singleplex).**

| | Primary PCR | | Nested PCR | |
|---|---|---|---|---|
| **Reagent** | **Final concentration** | **Volume per reaction** | **Final concentration** | **Volume per reaction** |
| $ddH_2O$ | - | 13.75 µL | - | 15.25 µL |
| Buffer B2 | 1x | 2.5 µL | 1x | 2.5 µL |
| dNTPs | 0.2 mM | 2.5 µL | 0.2 mM | 2.5 µL |
| $MgCl_2$ | 1.5 mM | 2.5 µL | 1.5 mM | 2.5 µL |
| Primers fw+rv | 400 nM | 1 µL | 400 nM | 1 µL |
| HOT FIREPol DNA polymerase ® | 0.05 U/µL | 0.25 µL | 0.05 U/µL | 0.25 µL |
| DNA | - | 2.5 µL | - | 1 µL |
| **Total volume** | - | 25 µL | - | 25 µL |

*Primary PCR products are diluted 1:10 in $ddH_2O$ before being added to nested PCR reaction.

**Supplementary Table 3. Primer sequences for *PfPK2, Poly-α, TA40* and *TA81*.**

| Primer | Primary PCR | Nested PCR |
|---|---|---|
| PfPK2_fw | 5'-CTTTCATCGATACTACGA-3' | 5'-CTTTCATCGATACTACGA-3' |
| PfPK2_rv | 5'-CCTCAGACTGAAATGCAT-3' | 5'-HEX-AAAGAAGGAACAAGCAGA-3' |
| Poly-α_fw | 5'-AAAATATAGACGAACAGA-3' | 5'-AAAATATAGACGAACAGA-3' |
| Poly-α_rv | 5'-ATCAGATAATTGTTGGTA-3' | 5'-FAM-GAAATTATAACTCTACCA-3' |
| TA40_fw | 5'-AAGGGATTGCTGCAAGGT-3' | 5'-AGGTAATAATAACACGAAC-3' |
| TA40_rv | 5'-GAAATTGGCACCACCACA-3' | 5'-PET-CATCAATAAAATCACTACTA-3' |
| TA81_fw | 5'-GAAGAAATAAGGGAAGGT-3' | 5'-FAM-TGGACAAATGGGAAAGGATA-3' |
| TA81_rv | 5'-TTTCACACAACACAGGATT-3' | 5'-TTTCACACAACACAGGATT-3' |

**Supplementary Table 4. Cycling conditions.**

| Primary PCR | | | Nested PCR | | |
|---|---|---|---|---|---|
| **Temperature** | **Time** | **Cycles** | **Temperature** | **Time** | **Cycles** |
| 95°C | 15:00 min | | 95°C | 15:00 min | |

| | | | | | |
|---|---|---|---|---|---|
| 95°C | 00:30 min | 30 | 95°C | 00:20 min | 25 |
| 45°C | 00:30 min | | 45°C | 00:20 min | |
| 65°C | 00:40 min | | 65°C | 00:30min | |
| 65°C | 05:00 min | | 65°C | 05:00 min | |
| 4°C | ∞ | | 4°C | ∞ | |

**Supplementary Table 5. Strain mixture samples.**

The longest amplicon for each marker is colored in red. The strain mixtures in which the longest amplicon is in minority are **bold**.

| | ***TA81*** | ***Poly-α*** | ***PfPK2*** | ***TA40*** |
|---|---|---|---|---|
| **Sample** | Ratio<br>3D7:K1:HB3:FCB1<br>119:125:128:116 | Ratio<br>3D7:K1:HB3:FCB1<br>152:177:180:152 | Ratio<br>3D7:K1:HB3:FCB1<br>168:168:191:161 | Ratio<br>3D7:K1:HB3:FCB1<br>207:186:192:186 |
| S5 | 1:1:1:1 | 1:1:1:1 | 1:1:1:1 | 1:1:1:1 |
| S6 | **5:1:1:1** | **5:1:1:1** | **5:1:1:1** | 5:1:1:1 |
| S7 | **1:5:1:5** | **1:5:1:5** | **1:5:1:5** | **1:5:1:5** |
| S8 | 1:5:5:5 | 1:5:5:5 | 1:5:5:5 | **1:5:5:5** |
| S9 | **1:10:1:1** | **1:10:1:1** | **1:10:1:1** | **1:10:1:1** |
| S10 | 1:1:10:10 | 1:1:10:10 | 1:1:10:10 | **1:1:10:10** |
| S11 | 10:1:10:10 | 10:1:10:10 | 10:1:10:10 | 10:1:10:10 |
| S12 | 1:1:15:1 | 1:1:15:1 | 1:1:15:1 | **1:1:15:1** |
| S13 | **15:15:1:1** | **15:15:1:1** | **15:15:1:1** | 15:15:1:1 |
| S14 | **15:15:1:15** | **15:15:1:15** | **15:15:1:15** | 15:15:1:15 |
| S15 | **1:1:1:20** | **1:1:1:20** | **1:1:1:20** | **1:1:1:20** |
| S16 | 20:1:20:1 | 20:1:20:1 | 20:1:20:1 | 20:1:20:1 |
| S17 | 20:20:20:1 | 20:20:20:1 | 20:20:20:1 | 20:20:20:1 |
| S18 | **50:1:1:1** | **50:1:1:1** | **50:1:1:1** | 50:1:1:1 |
| S19 | **1:50:1:50** | **1:50:1:50** | **1:50:1:50** | **1:50:1:50** |
| S20 | 1:50:50:50 | 1:50:50:50 | 1:50:50:50 | **1:50:50:50** |
| S21 | **100:1:1:1** | **100:1:1:1** | **100:1:1:1** | 100:1:1:1 |
| S22 | **1:100:1:100** | **1:100:1:100** | **1:100:1:100** | **1:100:1:100** |
| S23 | 1:100:100:100 | 1:100:100:100 | 1:100:100:100 | **1:100:100:100** |
| S27 | 1:1:1000:1 | 1:1:1000:1 | 1:1:1000:1 | **1:1:1000:1** |
| S28 | 1000:1000:1:1 | 1000:1000:1:1 | 1000:1000:1:1 | 1000:1000:1:1 |
| S29 | **1000:1000:1:1000** | **1000:1000:1:1000** | **1000:1000:1:1000** | 1000:1000:1:1000 |

**Supplementary Table 6. Samples list**.

| **Sample ID** | **Day** | **Study site** | **Comments** |
|---|---|---|---|
| 01/03D0 | 0 | Rukara | Excluded from R vs. NI |
| 01/03D19 | 19 | Rukara | Excluded from R vs. NI |
| 01/08D0 | 0 | Rukara | |
| 01/08D28 | 28 | Rukara | |
| 01/10D0 | 0 | Rukara | |
| 01/10D4 | 4 | Rukara | |
| 01/14D0 | 0 | Rukara | |
| 01/14D28 | 28 | Rukara | |
| 01/16D0 | 0 | Rukara | |

| | | | |
|---|---|---|---|
| 01/16D21 | 21 | Rukara | |
| 01/19D0 | 0 | Rukara | |
| 01/19D28 | 28 | Rukara | |
| 01/20D0 | 0 | Rukara | |
| 01/20D21 | 21 | Rukara | |
| 01/21D0 | 0 | Rukara | |
| 01/21D21 | 21 | Rukara | |
| 01/22D0 | 0 | Rukara | |
| 01/22D28 | 28 | Rukara | |
| 01/24D0 | 0 | Rukara | |
| 01/24D21 | 21 | Rukara | |
| 01/28D0 | 0 | Rukara | |
| 01/28D21 | 21 | Rukara | |
| 01/34D0 | 0 | Rukara | |
| 01/34D21 | 21 | Rukara | |
| 01/37D0 | 0 | Rukara | |
| 01/37D28 | 28 | Rukara | qPCR negative |
| 01/38D0 | 0 | Rukara | |
| 01/38D27 | 27 | Rukara | |
| 01/45D0 | 0 | Rukara | |
| 01/45D28 | 28 | Rukara | |
| 01/46D0 | 0 | Rukara | |
| 01/46D21 | 21 | Rukara | |
| 01/47D0 | 0 | Rukara | |
| 01/47D21 | 21 | Rukara | |
| 01/54D0 | 0 | Rukara | |
| 01/54D21 | 21 | Rukara | |
| 01/57D0 | 0 | Rukara | |
| 01/57D21 | 21 | Rukara | |
| 01/64D0 | 0 | Rukara | |
| 01/64D14 | 14 | Rukara | qPCR negative |
| 01/70D0 | 0 | Rukara | |
| 01/70D21 | 21 | Rukara | |
| 01/76D0 | 0 | Rukara | qPCR negative |
| 01/76D21 | 21 | Rukara | |
| 01/85D0 | 0 | Rukara | |
| 01/85D21 | 21 | Rukara | |
| 02/25D0 | 0 | Masaka | |
| 02/25D28 | 28 | Masaka | |
| 02/30D0 | 0 | Masaka | |
| 02/30D21 | 21 | Masaka | |
| 03/01D0 | 0 | Bugarama | |
| 03/01D21 | 21 | Bugarama | |
| 03/02D0 | 0 | Bugarama | |
| 03/02D21 | 21 | Bugarama | |
| 03/19D0 | 0 | Bugarama | |
| 03/19D28 | 28 | Bugarama | |
| 03/23D0 | 0 | Bugarama | |
| 03/23D21 | 21 | Bugarama | |
| 03/31D0 | 0 | Bugarama | |
| 03/31D14 | 14 | Bugarama | |
| 03/32D0 | 0 | Bugarama | |
| 03/32D28 | 28 | Bugarama | |
| 03/37D0 | 0 | Bugarama | |
| 03/37D21 | 21 | Bugarama | |
| 03/45D0 | 0 | Bugarama | |
| 03/45D15 | 15 | Bugarama | qPCR negative |
| 03/47D0 | 0 | Bugarama | |

| | | | |
|---|---|---|---|
| 03/47D21 | 21 | Bugarama | |
| 03/50D0 | 0 | Bugarama | |
| 03/50D28 | 28 | Bugarama | |
| 03/52D0 | 0 | Bugarama | |
| 03/52D28 | 28 | Bugarama | |
| 03/57D0 | 0 | Bugarama | |
| 03/57D21 | 21 | Bugarama | |
| 03/62D0 | 0 | Bugarama | |
| 03/62D27 | 27 | Bugarama | |
| 03/70D0 | 0 | Bugarama | |
| 03/70D21 | 21 | Bugarama | qPCR negative |
| 03/86D0 | 0 | Bugarama | |
| 03/86D28 | 28 | Bugarama | |
| 01/01D0 | 0 | Rukara | |
| 01/02D0 | 0 | Rukara | |
| 01/04D0 | 0 | Rukara | |
| 01/05D0 | 0 | Rukara | |
| 01/06D0 | 0 | Rukara | |
| 01/07D0 | 0 | Rukara | |
| 01/09D0 | 0 | Rukara | |
| 01/11D0 | 0 | Rukara | |
| 01/12D0 | 0 | Rukara | qPCR negative |
| 01/13D0 | 0 | Rukara | |
| 01/15D0 | 0 | Rukara | |
| 01/17D0 | 0 | Rukara | |
| 01/18D0 | 0 | Rukara | |
| 01/23D0 | 0 | Rukara | |
| 01/25D0 | 0 | Rukara | |
| 01/26D0 | 0 | Rukara | |
| 01/27D0 | 0 | Rukara | |
| 01/29D0 | 0 | Rukara | |
| 01/30D0 | 0 | Rukara | |
| 01/31D0 | 0 | Rukara | |
| 01/32D0 | 0 | Rukara | |
| 01/33D0 | 0 | Rukara | |
| 01/35D0 | 0 | Rukara | |
| 01/36D0 | 0 | Rukara | |
| 01/40D0 | 0 | Rukara | |
| 01/41D0 | 0 | Rukara | |
| 01/42D0 | 0 | Rukara | |
| 01/43D0 | 0 | Rukara | |
| 01/44D0 | 0 | Rukara | |
| 01/48D0 | 0 | Rukara | |
| 01/49D0 | 0 | Rukara | |
| 01/50D0 | 0 | Rukara | |
| 01/51D0 | 0 | Rukara | |
| 01/52D0 | 0 | Rukara | |
| 01/53D0 | 0 | Rukara | |
| 01/55D0 | 0 | Rukara | |
| 01/56D0 | 0 | Rukara | |
| 01/58D0 | 0 | Rukara | |
| 01/59D0 | 0 | Rukara | |
| 01/60D0 | 0 | Rukara | |
| 01/61D0 | 0 | Rukara | |
| 01/62D0 | 0 | Rukara | |
| 01/63D0 | 0 | Rukara | |
| 01/66D0 | 0 | Rukara | |
| 01/67D0 | 0 | Rukara | |

| | | | |
|---|---|---|---|
| 01/68D0 | 0 | Rukara | |
| 01/69D0 | 0 | Rukara | |
| 01/71D0 | 0 | Rukara | |
| 01/72D0 | 0 | Rukara | |
| 01/73D0 | 0 | Rukara | |
| 01/75D0 | 0 | Rukara | |
| 01/77D0 | 0 | Rukara | |
| 01/78D0 | 0 | Rukara | qPCR negative |
| 01/79D0 | 0 | Rukara | |
| 01/80D0 | 0 | Rukara | qPCR negative |
| 01/81D0 | 0 | Rukara | qPCR negative |
| 01/82D0 | 0 | Rukara | |
| 01/83D0 | 0 | Rukara | |
| 01/84D0 | 0 | Rukara | |
| 01/86D0 | 0 | Rukara | |
| 01/87D0 | 0 | Rukara | |
| 01/88D0 | 0 | Rukara | qPCR negative |
| 02/01D0 | 0 | Masaka | |
| 02/02D0 | 0 | Masaka | |
| 02/03D0 | 0 | Masaka | |
| 02/04D0 | 0 | Masaka | |
| 02/05D0 | 0 | Masaka | |
| 02/06D0 | 0 | Masaka | |
| 02/07D0 | 0 | Masaka | |
| 02/09D0 | 0 | Masaka | |
| 02/10D0 | 0 | Masaka | |
| 02/13D0 | 0 | Masaka | |
| 02/15D0 | 0 | Masaka | |
| 02/16D0 | 0 | Masaka | |
| 02/17D0 | 0 | Masaka | |
| 02/18D0 | 0 | Masaka | qPCR negative |
| 02/19D0 | 0 | Masaka | |
| 02/20D0 | 0 | Masaka | |
| 02/21D0 | 0 | Masaka | |
| 02/22D0 | 0 | Masaka | |
| 02/23D0 | 0 | Masaka | |
| 02/24D0 | 0 | Masaka | |
| 02/26D0 | 0 | Masaka | |
| 02/27D0 | 0 | Masaka | |
| 02/28D0 | 0 | Masaka | |
| 02/29D0 | 0 | Masaka | |
| 02/32D0 | 0 | Masaka | |
| 02/33D0 | 0 | Masaka | |
| 02/34D0 | 0 | Masaka | |
| 02/35D0 | 0 | Masaka | |
| 02/36D0 | 0 | Masaka | |
| 02/37D0 | 0 | Masaka | |
| 02/38D0 | 0 | Masaka | |
| 03/03D0 | 0 | Bugarama | |
| 03/03D2 | 2 | Bugarama | |
| 03/03D3 | 3 | Bugarama | |
| 03/04D0 | 0 | Bugarama | |
| 03/04D2 | 2 | Bugarama | |
| 03/04D3 | 3 | Bugarama | qPCR negative |
| 03/05D0 | 0 | Bugarama | |
| 03/05D2 | 2 | Bugarama | qPCR negative |
| 03/06D0 | 0 | Bugarama | |
| 03/07D0 | 0 | Bugarama | |

| | | | |
|---|---|---|---|
| 03/08D0 | 0 | Bugarama | |
| 03/09D0 | 0 | Bugarama | |
| 03/10D0 | 0 | Bugarama | |
| 03/11D0 | 0 | Bugarama | |
| 03/12D0 | 0 | Bugarama | |
| 03/13D0 | 0 | Bugarama | |
| 03/14D0 | 0 | Bugarama | |
| 03/15D0 | 0 | Bugarama | qPCR negative |
| 03/16D0 | 0 | Bugarama | |
| 03/17D0 | 0 | Bugarama | |
| 03/18D0 | 0 | Bugarama | |
| 03/20D0 | 0 | Bugarama | |
| 03/21D0 | 0 | Bugarama | |
| 03/22D0 | 0 | Bugarama | |
| 03/24D0 | 0 | Bugarama | |
| 03/25D0 | 0 | Bugarama | |
| 03/26D0 | 0 | Bugarama | |
| 03/27D0 | 0 | Bugarama | |
| 03/29D0 | 0 | Bugarama | |
| 03/33D0 | 0 | Bugarama | |
| 03/34D0 | 0 | Bugarama | |
| 03/35D0 | 0 | Bugarama | |
| 03/38D0 | 0 | Bugarama | |
| 03/39D0 | 0 | Bugarama | |
| 03/40D0 | 0 | Bugarama | |
| 03/41D0 | 0 | Bugarama | |
| 03/42D0 | 0 | Bugarama | |
| 03/43D0 | 0 | Bugarama | |
| 03/44D0 | 0 | Bugarama | |
| 03/46D0 | 0 | Bugarama | |
| 03/48D0 | 0 | Bugarama | |
| 03/49D0 | 0 | Bugarama | qPCR negative |
| 03/53D0 | 0 | Bugarama | |
| 03/54D0 | 0 | Bugarama | |
| 03/55D0 | 0 | Bugarama | |
| 03/56D0 | 0 | Bugarama | |
| 03/58D0 | 0 | Bugarama | |
| 03/59D0 | 0 | Bugarama | |
| 03/60D0 | 0 | Bugarama | |
| 03/63D0 | 0 | Bugarama | |
| 03/64D0 | 0 | Bugarama | |
| 03/65D0 | 0 | Bugarama | |
| 03/66D0 | 0 | Bugarama | |
| 03/67D0 | 0 | Bugarama | |
| 03/68D0 | 0 | Bugarama | |
| 03/69D0 | 0 | Bugarama | |
| 03/71D0 | 0 | Bugarama | |
| 03/73D0 | 0 | Bugarama | |
| 03/74D0 | 0 | Bugarama | |
| 03/75D0 | 0 | Bugarama | |
| 03/76D0 | 0 | Bugarama | |
| 03/77D0 | 0 | Bugarama | Insufficient blood on DBS |
| 03/78D0 | 0 | Bugarama | |
| 03/79D0 | 0 | Bugarama | |
| 03/80D0 | 0 | Bugarama | |
| 03/81D0 | 0 | Bugarama | |
| 03/82D0 | 0 | Bugarama | qPCR negative |
| 03/83D0 | 0 | Bugarama | |

| | | |
|---|---|---|
| 03/84D0 | 0 | Bugarama |
| 03/85D0 | 0 | Bugarama |
| 03/88D0 | 0 | Bugarama |
| 02/39D0 | 0 | Masaka |
| 02/40D0 | 0 | Masaka |
| 01/39D0 | 0 | Rukara |
| 01/65D0 | 0 | Rukara |
| 01/74D0 | 0 | Rukara |
| 02/41D0 | 0 | Masaka |
| 02/42D0 | 0 | Masaka |
| 02/08D0 | 0 | Masaka |
| 02/11D0 | 0 | Masaka |
| 02/12D0 | 0 | Masaka |
| 02/14D0 | 0 | Masaka |
| 02/31D0 | 0 | Masaka |
| 03/28D0 | 0 | Bugarama |
| 03/30D0 | 0 | Bugarama |
| 03/36D0 | 0 | Bugarama |
| 03/51D0 | 0 | Bugarama |
| 03/61D0 | 0 | Bugarama |
| 03/72D0 | 0 | Bugarama |
| 03/87D0 | 0 | Bugarama |
| 02/43D0 | 0 | Masaka |
| 02/44D0 | 0 | Masaka |
| 02/45D0 | 0 | Masaka |
| 02/46D0 | 0 | Masaka |
| 02/47D0 | 0 | Masaka |
| 02/48D0 | 0 | Masaka |
| 02/49D0 | 0 | Masaka |
| 02/50D0 | 0 | Masaka |
| 02/51D0 | 0 | Masaka |
| 02/52D0 | 0 | Masaka |

**Supplementary Table 7. Samples with discordant recrudescence (R) and new infection (NI) results depending on marker combination. Results by single marker.**

Allelic families (K1, MAD20, Ro33 for *msp1* and 3D7, FC27 for *msp2*) are indicated in blue.

| Sample ID | K1 | MAD20 | Ro33 | *msp1* | 3D7 | FC27 | *msp2* | *glurp* | *Poly-α* | *TA81* | *TA40* | *PfPK2* |
|---|---|---|---|---|---|---|---|---|---|---|---|---|
| 01/14 | NI | NI | R | R | NI | R | R | R | NI | R | R | R |
| 01/28 | NI | NI | R | R | NI | R | R | R | NI | NI | NI | R |
| 01/47 | NI | NI | R | R | R | NI | R | NI | R | NI | NI | R |
| 03/02 | NI | NI | R | R | NI | R | R | NI | R | R | R | R |

**Supplementary Table 8. Samples with discordant recrudescence (R) and new infection (NI) results depending on the method used. Results by single marker.**

Allelic families (K1, MAD20, Ro33 for *msp1* and 3D7, FC27 for *msp2*) are indicated in blue.

| Sample ID | *ama1-D2* | *ama1-D3* | *cpmp* | K1 | MAD20 | Ro33 | *msp1* | 3D7 | FC27 | *msp2* | *glurp* | 7 micro-satellites (prob. of R) |
|---|---|---|---|---|---|---|---|---|---|---|---|---|
| 01/14 | NI | NI | NI | NI | NI | R | R | NI | R | R | R | NI (0.003) |
| 01/21 | R | R | R | NI | NI | R | R | NI | R | R | R | NI (0.001) |
| 01/28 | R | R | NI | NI | NI | R | R | NI | R | R | R | NI (0.000) |
| 01/54 | NI | NI | NI | NI | NI | NI | NI | NI | NI | NI | NI | R (0.525) |
| 03/02 | R | R | R | NI | NI | R | R | NI | R | R | NI | R (0.857) |

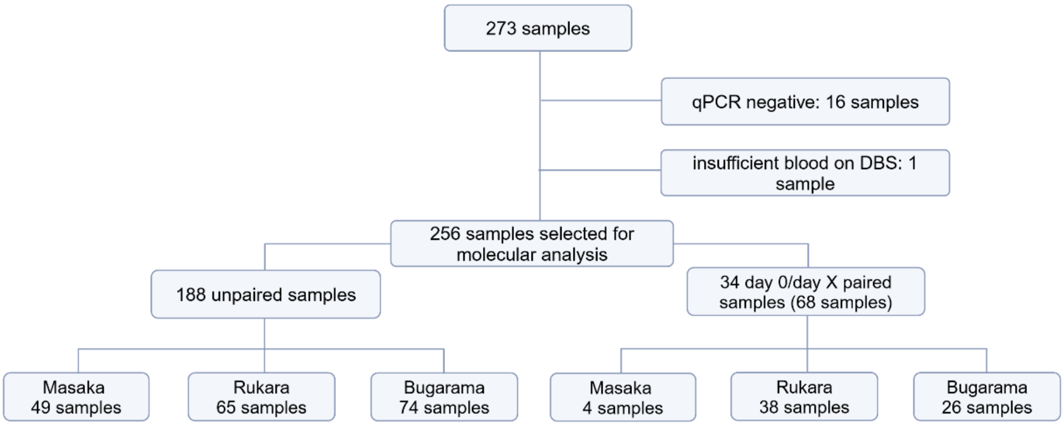

**Supplementary Figure 1. Type, origin and number of samples in analysis.**

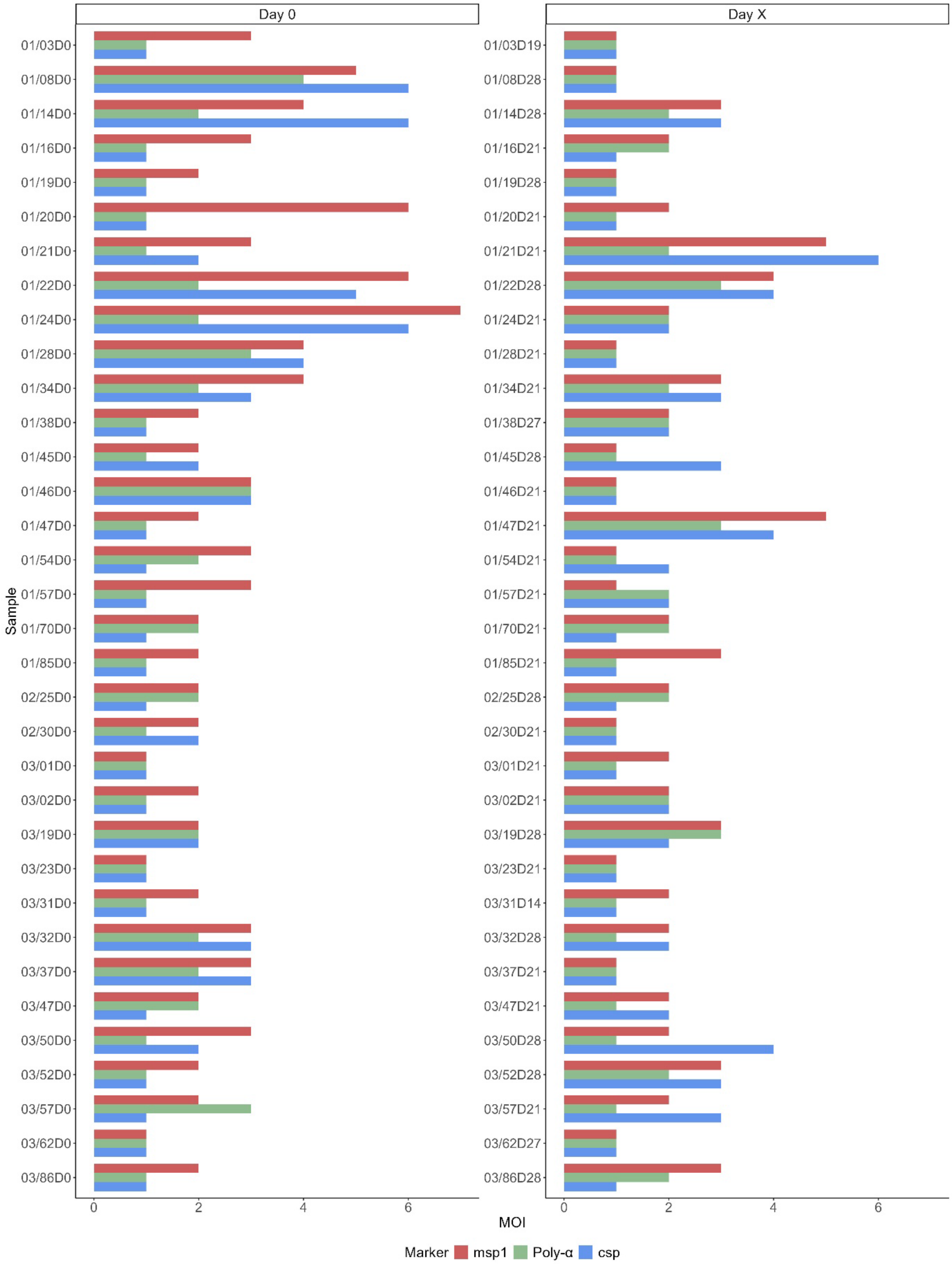

**Supplementary Figure 2. MOI value detected by *msp1, Poly-α* and *csp* in Day 0 and Day X samples.**
